# Epididymal extracellular vesicles harbor and convey mRNA to sperm for transfer to zygotes

**DOI:** 10.1101/2025.08.18.670952

**Authors:** Natalie A. Trigg, Grace S. Lee, Alexis G. Leach, Colin C. Conine

## Abstract

The epididymis plays a critical role in sperm maturation, including remodeling the sperm RNA payload. While small RNAs have been extensively studied in this context, the epididymal contribution to larger sperm RNAs, such as mRNAs, remains underexplored. mRNAs were among the first RNA species identified in sperm, yet their functional relevance has remained elusive, largely due to the translational quiescence of mature spermatozoa and the hypothesis that these RNAs are residual by-products of spermatogenesis. Yet, mRNAs carried by sperm have been detected in the zygote, indicating they could play a role beyond fertilization. However, if the soma, the epididymis, actively contributes mRNAs to sperm as it does small RNAs, has not been experimentally assessed. To investigate this, here, we provide a comprehensive analysis of the mRNA landscape of mouse sperm, epithelial cells and extracellular vesicles (EVs) isolated from the proximal (caput) and distal (cauda) epididymis. Through this analysis and sperm-EV co-incubation experiments we demonstrate the transfer of mRNAs from epididymal EVs to sperm. Further, through sperm RNA microinjection into zygotes we uncover gene regulation in the early embryo driven by the introduction of sperm RNAs, specific to large RNA species. These findings ultimately reveal the dynamic mRNA profile of sperm that is delivered to the egg and demonstrate that RNA species beyond small RNAs are capable of influencing preimplantation embryo gene expression.

## INTRODUCTION

Mature spermatozoa harbor a complex array of RNAs, including small non-coding RNAs (miRNA, piRNAs, tRNA-derived RNAs (tDRs or also known as tRFs) and long non-coding RNAs; lncRNAs) as well as coding RNAs, such as messenger RNAs (mRNAs) (Ostermeier et al., 2002, Krawetz et al., 2011, Peng et al., 2012, Sharma et al., 2016). During germ cell development, the RNAs present in sperm reflect the transcription required for sperm development. However, at the culmination of spermatogenesis, particularly the differentiation of haploid round spermatids to differentiated spermatozoa (spermiogenesis) genomic compaction precludes de novo RNA synthesis (Johnson et al., 2011, Miller, 2014). Thus, all RNAs present in sperm must be transcribed during spermatogenesis, processed from spermatogenic precursor transcripts, or acquired non-autonomously. It is widely thought that the majority of sperm mRNAs arise from transcription during spermatogenesis and remain in the differentiated spermatozoa. Indeed, highly abundant mRNAs detected in mature epididymal sperm are also detected at high abundance in germ cells during spermatogenesis in the testis (i.e. *Protamine 1* and *2*) (Miller, 2000, Djureinovic et al., 2014, Sendler et al., 2013). Numerous studies have detected a rich profile of mRNA transcripts within the sperm of multiple species, including humans and mice (Sun et al., 2021, Ostermeier et al., 2002). However, whether these RNAs are, as thought, simply the remnants of the spermatogenic gene expression program or if they have functions post-fertilization in the zygote is not thoroughly understood. Post-fertilization functions of sperm RNAs have been proposed, for example, sperm *Igf2* mRNA expression correlates with embryo morphokinetics in humans, which is supported by a recent study that examined the mechanistic basis of this correlation, by microinjecting synthesized *Igf2* mRNA into mouse parthenotes and assessing gene expression (Cannarella et al., 2024). Microinjection of *Igf2* mRNA led to altered expression of 99 genes in the early embryo. Similarly, other sperm-delivered mRNAs, such as *Dby* have been implicated in zygotic development through microinjection of antisense RNA to *Dby* mRNA, which prohibits embryonic development to the 2-cell stage in mice (Yao et al., 2010). Moreover, sperm RNAs greater than 200 nucleotides have been shown to be modulated by paternal stress exposure and contribute to specific non-genetically inherited trauma symptoms in offspring, including food intake and glucose response to insulin (Gapp et al., 2020). Such findings highlight a functional role for sperm delivered mRNAs in the embryo and uncover another layer of non-genetic information harbored by the sperm cell.

The accumulation of mature sperm mRNAs outside of spermatogenic persistence, such as through post-transcriptional processing or non-autonomous accumulation in the epididymis has yet to be thoroughly addressed. While certain types of sperm RNAs could be processed from spermatogenic precursors in the epididymis, such as small RNAs, the de novo generation of mRNAs in epididymal sperm is unlikely as they are typically processed co-transcriptionally (Bentley, 2014). However, the acquisition of mRNAs during epididymal transit from the epididymal epithelium is plausible, as this mechanism has been clearly demonstrated to occur for small RNAs (Trigg and Conine, 2024, Sharma et al., 2018, Reilly et al., 2016). The epididymis is a long-coiled tubule, positioned adjacent to the testis and facilitates sperm maturation following sperm development in the testis. During transit through the epididymis, sperm acquire motility and undergo membrane remodeling that renders them capable of fertilization (Weigel Muñoz et al., 2024). Beyond these functional gains, sperm also acquire macromolecules during epididymal transit. A notable example is the increased abundance of 57 sperm proteins, including carbonic anhydrase 4 (CA4) and calcium-binding tyrosine phosphorylation-regulated protein (CAYBR) and post translational modifications to existing proteins (Sharma et al., 2018, Reilly et al., 2016, Martin-DeLeon, 2015, Skerrett-Byrne et al., 2022). Additionally, it has been shown that the small RNA profile of sperm is dramatically remodeled as the sperm passage from the proximal (caput) epididymis to the distal (cauda) epididymis (Nixon et al., 2015b, Sharma et al., 2018). Specifically, a subset of miRNAs are transferred to sperm during the transition from the caput to cauda epididymis and, importantly, these miRNAs were demonstrated to regulate preimplantation embryonic gene expression (Conine et al., 2018, Trigg and Conine, 2024, Trigg et al., 2019). Interestingly, miRNAs and tDRs are delivered to maturing sperm by first being packaged into extracellular vesicles, called epididymosomes, within the epididymal epithelium (Reilly et al., 2016, Sharma et al., 2018).

Although the mRNA profile of epididymal sperm has not previously been comprehensively assessed across the epididymis, evidence suggests that, like other macromolecules, it may change during transit, with transcripts potentially gained and lost between the caput and cauda (Robertson et al., 2020). This challenges the notion that sperm mRNAs are merely remnants of testicular development and suggests that a mechanism exists for epididymal sperm to acquire and lose mRNAs. If packaged with mRNAs, the EVs produced by the epididymal epithelium could provide a mechanism of mRNA acquisition by epididymal sperm. Moreover, the presence of big RNAs (>200 nucleotides), likely to include mRNAs, lncRNAs and circular RNAs (circRNAs), have been observed in epididymal EVs using gel electrophoresis (Conine et al., 2018). These findings led us to hypothesize that extracellular vesicles deliver mRNA to sperm during epididymal transit. To assess this, we investigated the mRNA profile of extracellular vesicles, epithelial cells and sperm isolated from the caput and cauda epididymis via mRNA-seq. In extending this analysis to understand the contribution of EVs to the sperm mRNA profile we also performed co-incubation experiments to demonstrate that EVs transfer mRNA to sperm *in vitro*. Lastly, to begin uncovering the functional role of sperm RNAs longer than 200 nucleotides, we used parthenogenetically activated eggs combined with microinjection to evaluate how different fractions of sperm RNA influence embryonic gene expression.

## MATERIALS AND METHODS

### Animals

Adult FVB/NJ male mice (8-12 weeks old) and female mice (5-7 weeks old) were derived from mice purchased from Jackson Labs (Strain #:001800) and breed in-house. All animals used in this study were handled and kept in conditions under strict compliance with the Children’s Hospital of Philadelphia and the University of Pennsylvania Institutional Animal Care and Use Committee regulations (CHOP IACUC Protocol #23-001364, UPenn PSOM IACUC Protocol #806911)

### Epididymal sperm and extracellular vesicle isolation

Epididymides were dissected from male mice and separated into the proximal caput epididymis (including the initial segment) and the distal cauda epididymis (Fig. S1B).

Dissected epididymal tissue was placed in a dish containing 1.5 mL of TYH media (120 mM NaCl, 4.7 mM KCl, 1.7 mM CaCl_2_2H_2_O, 1.2 mM KH_2_PO_4_, 1.2 mM MgSO_4_, 25 mM NaHCO_3_, 5.5 mM glucose, 0.5 mM sodium pyruvate, 10 µg/mL gentamicin, 3.0 mg/mL bovine serum albumin and 1.0% phenol red). The luminal contents of the cauda epididymis were gently squeezed into the media under a dissection microscope being careful to minimize tissue disruption. For the caput epididymis, small incisions were made into the tissue to release the luminal fluid into the media. Tissue was removed from the dish and sperm was allowed to disperse for 10 min at 37°C. For cauda sperm isolation, after incubation the media was transferred to a microcentrifuge tube and sperm were allowed to swim up for 10 min at 37°C, before filtering over a 70 µm filter and transferred to a new tube and centrifuged 5 min at 500 × *g* for 5 min. Following this centrifugation, the supernatant was transferred to a new tube and subjected to differential centrifugation to isolate extracellular vesicles (EVs) as previously described (Trigg and Conine, 2024). Sperm pellets were retained to purify populations of sperm. Cauda sperm pellets were washed in phosphate buffered saline (PBS), incubated in somatic cell lysis buffer on ice for 10 min and subsequently washed prior to snap freezing. Sperm washes following somatic cell lysis was performed at 10,000 × *g* at 4°C to pellet all cells and minimize cell loss. Caput sperm pellets were resuspended in 1 mL of media following removal of supernatant and layered over a 28% Percoll gradient and centrifuged for 15 min, 400 × *g* at 37°C. Pelleted sperm were washed in PBS and any remaining somatic cells were removed by incubation in somatic cell lysis buffer for 10 min on ice. Sperm were washed a final time in PBS and snap frozen. Incubation in somatic cell lysis buffer (0.01% SDS, 0.05% Triton-X-100) also removed adherent cytoplasmic droplets from spermatozoa, which ensured we assayed the population of RNAs in mature sperm (Fig. S1D-E).

Extracellular vesicle isolation was confirmed using particle tracking (Fig. S2A) and negative stain electron microscopy. Particle analysis was performed using the Particle Metric ZetaView and negative stain electron microscopy was conducted by the University of Pennsylvania Electron Microscopy Resource Lab (RRID:SCR_022375).

### RNase treatment of extracellular vesicles

Extracellular vesicles were resuspended in 300μl of cold PBS and divided equally among treatments. Samples were then incubated in RNase A for 20 min at 37°C at a concentration 0.5 µg/mL in PBS to eliminate any unprotected RNA. RNase A activity was inactivated via incubation at-80°C for 5 min and then vesicle preparations were immediately homogenized in TRI-reagent and total RNA was extracted. Mock treatments were incubated as above with the addition of vehicle only (PBS) at equal volumes in place of RNase A.

### Sperm RNA extraction and preparation for microinjection

RNA was extracted from populations of mature spermatozoa collected from 20 mice as previously described (Trigg and Conine, 2024). Total RNA was quantitated, and a portion of total RNA was diluted and aliquoted for injection. The remaining RNA was subjected to size selection (>200 nucleotides) using the Monarch Total RNA Miniprep kit (NEB, T2010S). After size selection of the RNA populations, RNA was diluted to desired concentration and H3.3.-GFP mRNA was added ready for microinjection.

### Epididymal epithelial cell purification

Epithelial cells were purified from caput and cauda epididymal tissue as previously described (Trigg et al., 2022, Nixon et al., 2015a, Trigg et al., 2021). Populations of epithelial cells were checked for purity by staining with nuclear stain, DAPI (4′,6-diamidino-2-phenylindole) and recording the percentage of somatic cell purity. Only samples with >80% purity were kept for RNA-seq.

### Sperm-extracellular vesicle co-incubation

Cauda EVs were isolated as outlined above and resuspended in 50 µL of TYH media, mixed well to resuspend vesicles into solution and kept at room temperature. Caput spermatozoa were isolated as described above with slight modification. Following incubation after tissue dispersion, the sperm containing supernatant was immediately filtered over a 70 µm cell strainer and then loaded onto the Percoll gradient. Following centrifugation, sperm was resuspended in TYH media and washed twice before resuspension in 200 µL media and split into two. Cauda EVs (50 µL) were added to one tube and 50 µL of media to the control and allowed to incubate for 2 h at 37°C with constant slow rocking. At the completion of the co-incubation 850 µL of PBS was added and sperm were pelleted for 5 min at 500 × *g*. Cells were incubated in somatic cell lysis buffer for 7 min and washed in PBS before snap freezing. Centrifugation following somatic cell lysis was performed at 10,000 × *g* for 5 min at 4°C.

### In vitro fertilization and zygote collection

Female 5–7-week-old FVB/NJ mice were superovulated by intraperitoneal injection of 5 IU pregnant mare serum gonadotropin (PMSG) and 5 IU of human chorionic gonadotropin (hCG) 48 h later. Cumulus-oocyte complexes (COCs) were retrieved from the distal oviductal ampullae 13-15 h after hCG injection and recovered in warmed PBS. COCs were then divided into two, where the first was added to a droplet of human tubal fluid (HTF) supplemented with 1.0 mM reduced glutathione (GSH) ready for in vitro fertilization (IVF) while the remaining COC’s were dissociated via brief incubation in hyaluronidase (3 mg/mL) and eggs washed of any residual cumulus cells and incubated in KSOM for 30 min under an atmosphere of 5% O_2_, 5% CO_2_. After equilibration, cumulus cell free eggs were collected into lysis buffer (1 x TCL buffer (Qiagen, Cat# 1070498) supplemented with 1% β-mercaptoethanol) and frozen in preparation for RNA-seq. Spermatozoa were collected from the cauda epididymis as described above and allowed to swim out in Biggers Whitten and Whittingham media (BWW; composed of 91.5 mM NaCl, 4.6 mM KCl, 1.7 mM CaCl_2_2H_2_O, 1.2 mM KH_2_PO_4_, 1.2 mM MgSO_4_7H_2_O, 25 mM NaHCO_3_, 5.6 mM D-glucose, 0.27 mM sodium pyruvate, 44 mM sodium lactate, 5 U/mL penicillin, 5 μg/mL streptomycin, 20 mM HEPES buffer, and 3.0 mg/mL bovine serum albumin [BSA]) containing 1.0 mg/mL of methyl-β-cyclodextrin) for 45 min at 37°C under an atmosphere of 5% O_2_, 5% CO_2_. Following capacitation, spermatozoa were added to the COC containing droplet and incubated for 3 h. After co-incubation, presumptive zygotes were thoroughly washed free of any bound spermatozoa and cultured in KSOM. To collect zygotes prior to the initiation of zygotic genome activation, 1 cell embryos were collected 4.5 h after fertilization, a time point when zygotic genome activation has yet to commence (Bouniol et al., 1995, Yang et al., 2016). Only zygotes with two pronuclei (fertilized) were collected. Zygotes were collected into lysis buffer and stored at-80°C for RNA-seq.

### Parthenogenesis, microinjection and embryo culture

Cumulus-oocyte complexes (COCs) from 12 superovulated female mice (per replicate) were collected as described above into a single dish of filtered PBS. COCs with an estimated egg count of ∼20 (typically 1-2 COC) were immediately transferred to a drop of supplemented HTF for IVF as described above. The remaining COCs were incubated in hyaluronidase to remove cumulus cells (see above). Following removal of cumulus cells, eggs were washed in KSOM before transfer to a droplet of KSOM supplemented with 10 mM SrCl_2_, 4 mM EGTA and cytochalasin B at a final concentration of 0.05 µg/mL to activate by incubation at 37°C under an atmosphere of 5% O_2_, 5% CO_2_ for 1 h (Ma et al., 2005). Following activation, parthenotes were washed through six droplets of KSOM ready for microinjection.

Parthenotes were transferred to the injection plate, in droplets of flushing and holding media (FHM) supplemented with 1% polyvinyl alcohol (PVA) for micromanipulation. Control injections consisted of H3.3.-GFP mRNA alone (50 ng/ul) and experimental injections included total RNA (100 ng/ul) or big RNA (50 ng/ul) isolated from populations of mature mouse spermatozoa. All experimental RNA injections also had H3.3-GFP mRNA added to injection mix at a final concentration of 50 ng/ul. RNA injections were carried out using a Femtojet (Eppendorf/Calibre) microinjector with Femtotip II microinject capillary tips at 100 hPa pressure for 0.2 s, with 7 hPa compensation pressure (Lee et al., 2025). After RNA injection, parthenotes were washed in KSOM and cultured. The presence of GFP signal was confirmed using fluorescence microscopy and any negative 2-cell embryos were removed from the culture dish. Injected embryos were cultured and collected for single-embryo RNA-sequencing. Four-cell and morula stage embryos were collected into lysis buffer at 46 and 72 h post-fertilization, respectively.

### RNA-sequencing and analysis

RNA was extracted from mouse sperm, epithelial cells and EVs as previously described (Trigg and Conine, 2024, Sharma et al., 2018, Trigg et al., 2025). Contaminating DNA was removed from total RNA samples by incubation with DNase I (Qiagen, 79254) as per the manufacturer’s instructions. RNA was quantified using Qubit fluorometer and 2 ng of RNA was used for RNA-seq library generation. For embryos, following lysis, RNA was isolated via RNAClean-XP beads (Beckman Coulter, Cat# A63987) as outlined previously (Harris et al., 2024). Libraries were generated for sequencing using the SMART-seq protocol as previously described (Trombetta et al., 2014, Trigg and Conine, 2024). Data were mapped to the *Mus musculus* reference genome (mm10) using Feature counts on ViaFoundry (Yukselen et al., 2020) and normalized to transcripts per million (TPM). Sequencing quality control was assessed by determining the number of transcripts within each sample with TPM ≥1. Sperm, epithelial cell and EV samples with less than 5,500 and egg/embryo samples with less than 8,000 genes satisfying this cutoff were removed. Raw read counts were uploaded to R Statistical Software and using the DESeq2 package data was normalized and differential expression of genes was determined using false-discovery rate *P*-value (Love et al., 2014). All datasets were further filtered to obtain a list of ‘expressed’ genes within each sample. This filtering involved removing any transcripts with less than 5 TPM average in the groups being compared. Except for embryo datasets, further filtering involved the removal of transcripts with less than half of replicates exhibiting TPM ≤ 2. Eulerr package in R was used to determine the overlapping genes among different groups and generate Venn diagrams and DEGs.

### Assessment of RNA conservation in human sperm

Human sperm low input mRNA-seq data was used to determine the conservation of sperm mRNAs between mice and humans. Utilizing the Ensembl (release #13) Bioconductor BiomaRt package for R, a list of all human orthologs to mouse genes was created. Gene common names used during sequence mapping were converted to Ensembl IDs using BiomaRt. (external gene name, external synonym, MGI symbol, and NCBI RefSeq ID were searched). During this conversion, 226 gene names could not be matched to an Ensembl ID using BiomaRt and were therefore excluded. Human orthologs were then called for each gene in the total list of genes mapped during sequencing. Abundance of human orthologs was determined and plotted against mouse gene abundance for the mRNAs detected in mature mouse sperm.

## RESULTS

### The mRNA profile of epididymal sperm, epithelial cells and extracellular vesicles is modified along the epididymis

To quantify the mRNA expression dynamics that occur in epididymal sperm, we performed mRNA-seq from mice, on caput (proximal) and cauda (distal) sperm, epididymal epithelial cells, and extracellular vesicles (EVs) isolated from epididymal fluid from mice. Importantly, these experiments confirm the presence of mRNAs within epididymal sperm total RNA preparations (Fig. S1A). While small RNA delivery from EVs to sperm during epididymal transit is well established (Conine et al., 2018, Sharma et al., 2018, Reilly et al., 2016), the role of EVs in transferring big RNAs, particularly mRNAs, remains unexplored. Further, we have previously shown abundant big RNA (> 200 nucleotides) species are present in EVs isolated from the caput and cauda epididymis via denaturing PAGE of total RNA (Conine et al., 2018). Thus, we hypothesize that epididymal EVs harbor mRNAs with dynamic accumulation along the epididymis that could deliver mRNAs to sperm.

Principal component analysis (PCA) grouped biological replicates together and separated samples by sample type and epididymal segment (Fig. 1A). Determination of Pearson correlations highlighted similar mRNA expression in sperm and EVs, while epithelial cells were distinct (Fig. S1C). Further, the mRNA profile of sperm, epithelial cells and vesicles isolated from the caput epididymis were distinctly separate from those isolated from the cauda epididymis. In exploring the dynamic changes that occur in mRNA profiles along the epididymis, we compared filtered gene lists of sperm, epithelial cells, and EVs from the caput and cauda epididymis. This comparison revealed that over 50% of transcripts in sperm and epithelial cells are present in both segments of the epididymis (Fig. 1B-C). Compared to sperm and epithelial cells, EVs from both the caput and cauda expressed significantly less mRNA species (Fig. 1D). Moreover, in EVs, only 33.7% of transcripts were shared in datasets from the caput and cauda. Rather, significantly more transcripts were detected in EVs isolated from the cauda (61.7% of mRNAs were unique to cauda) compared to the caput (4.6% unique to cauda; Fig. 1D). Examination of the five most abundant genes across sample type and epididymal segment highlighted segment specific genes (e.g. *Defb20* and *Crisp1*) and sample type specific genes (e.g. *Prm2* and *Flt1*; Fig. 1E). Next, we compared detected transcripts across sample types from the same segment to determine shared and unique genes (Fig. 1F-G). For both epididymal segments examined, sperm harbored the largest number of unique transcripts that were not detectable in either epithelial cells or EVs (>2,000 mRNAs unique to sperm). The percentage of transcripts expressed in all three sample types was comparable between the caput (24.2%) and cauda (31.2%). Moreover, EVs displayed the least number of unique transcripts of all sample types, with the majority of EV transcripts (98.0% in the caput and 89.3% in the cauda) also identified in other samples (Fig. 1F-G). Overall, this comparison highlighted three key findings, i) sperm are a specialized cell with a large number of unique transcripts that are not shared with epididymal epithelial cells or EVs and therefore likely remnants of germline transcription occurring at the culmination of spermatogenesis, ii) the EV mRNA profile is less complex than their parent cell (epithelial cells) and iii) RNA within EVs is also present in either sperm or epithelial cells.

**Figure 1:**
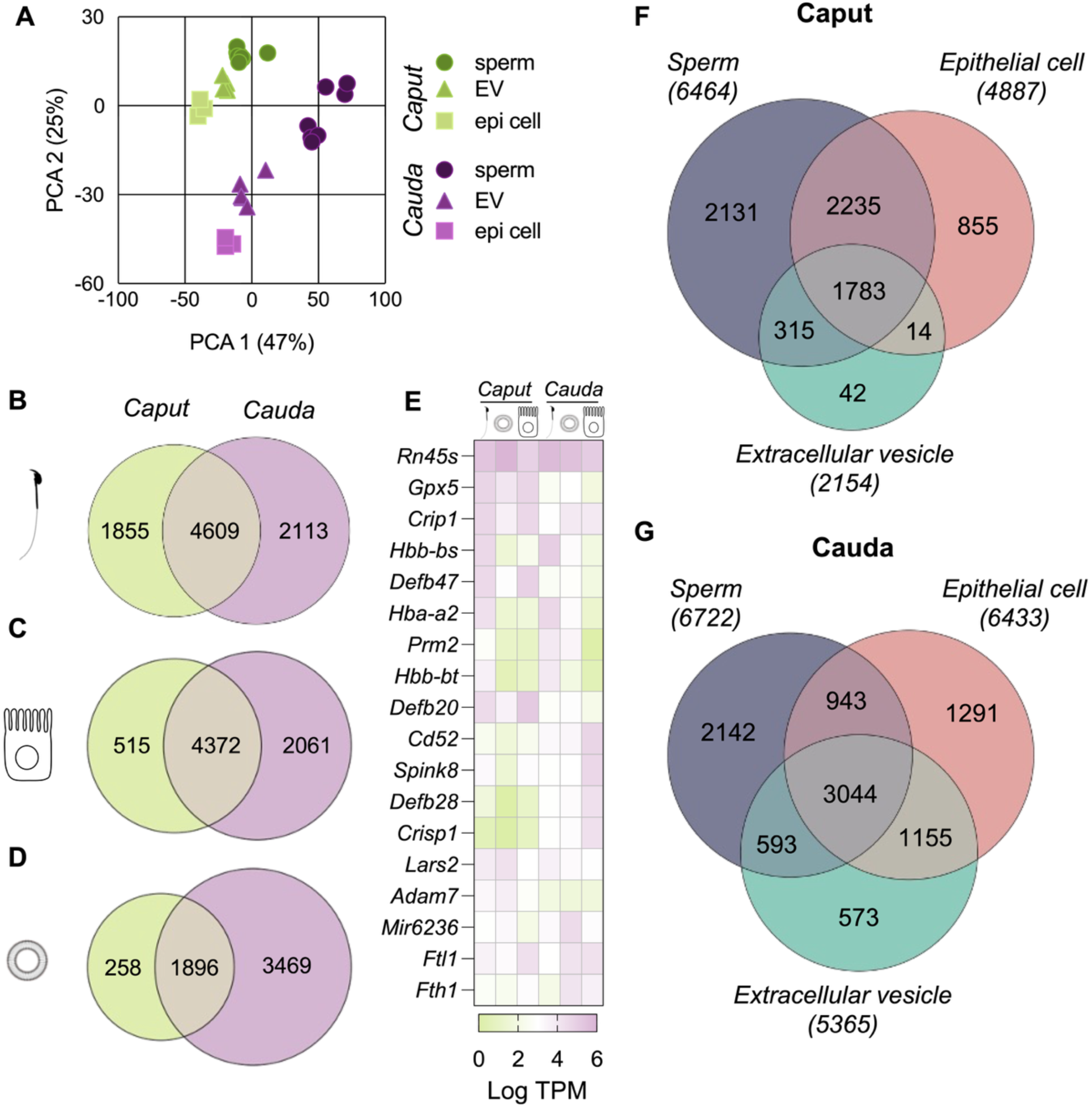
**Segment specific expression of epididymal mRNAs in sperm, epithelial cells and epididymosomes**. A) Principal component analysis (PCA) of the transcriptome of 38 individual samples of sperm, epithelial cells (epi cell) and epididymal extracellular vesicles (EVs) isolated from the caput (green) and cauda (purple) epididymis. A minimum of three biological replicates were sequenced (n= 3-8), with each biological replicate containing pooled samples from 3-4 mice. B-D) Proportional Venn diagram comparing the mRNA profile of genes expressed in B) sperm, C) epithelial cells and D) EVs isolated from the caput and cauda epididymis. E) Heatmap depicting the expression (log transcripts per million; TPM) of the top 5 most abundant genes across sample type and epididymal segment. F-G) Proportional Venn diagrams illustrating the number of identified genes expressed in populations of sperm, epithelial cells and EVs isolated from F) the caput and G) the cauda epididymis. Raw gene lists for each sample were filtered to determine the list of ‘detected’ genes which were compared here. Genes with no detectable expression (i.e., 0 TPM) in ≥ 50% of replicates were removed and the list was further filtered to remove genes with average TPM value of σ; 5 across all replicates.

Despite being transcriptionally silent, mature spermatozoa isolated from the distal cauda epididymis harbor many unique transcripts. Hence, we next sought to establish the dynamic changes in the RNA profile along the epididymis. Using DESeq2 we determined differentially expressed genes (DEGs) within sperm, epithelial cells and epididymosomes isolated from the caput and the cauda epididymis. This analysis revealed the differential expression of 2,417 genes in caput sperm compared to cauda sperm (Fig. 2A). Of these genes, 1,318 were more abundant in caput sperm, while the remaining 1,099 were detected at higher levels in cauda sperm (Fig. 2A). The expression profile of epithelial cells and EVs was also significantly altered along the epididymis. Epithelial cells isolated from the cauda epididymis displayed altered expression of 425 genes: 183 of which exhibited increased expression, while 242 decreased in expression compared to caput epithelial cells (Fig. 2B). In line with this, mRNA from EVs also differed depending on the epididymal location, with mRNAs from 289 genes significantly increased in cauda EVs compared to caput, while 304 were decreased (Fig. 2C). Comparison of DEGs across sample types revealed that while a portion of mRNAs were equally altered in all sample types from the caput to the cauda, the majority of DEGs were unique to each sample type (Fig. 2D-E, S1F-H). Despite this, correlating the caput to cauda fold change (log_2_) of mRNAs in sperm and EVs revealed a significant correlation (Pearson’s correlation coefficient of 0.33, Fig. 2F). These findings establish that the epididymal expression dynamics of many mRNAs in sperm are similar between sperm and EVs transiting the epididymis.

**Figure 2:**
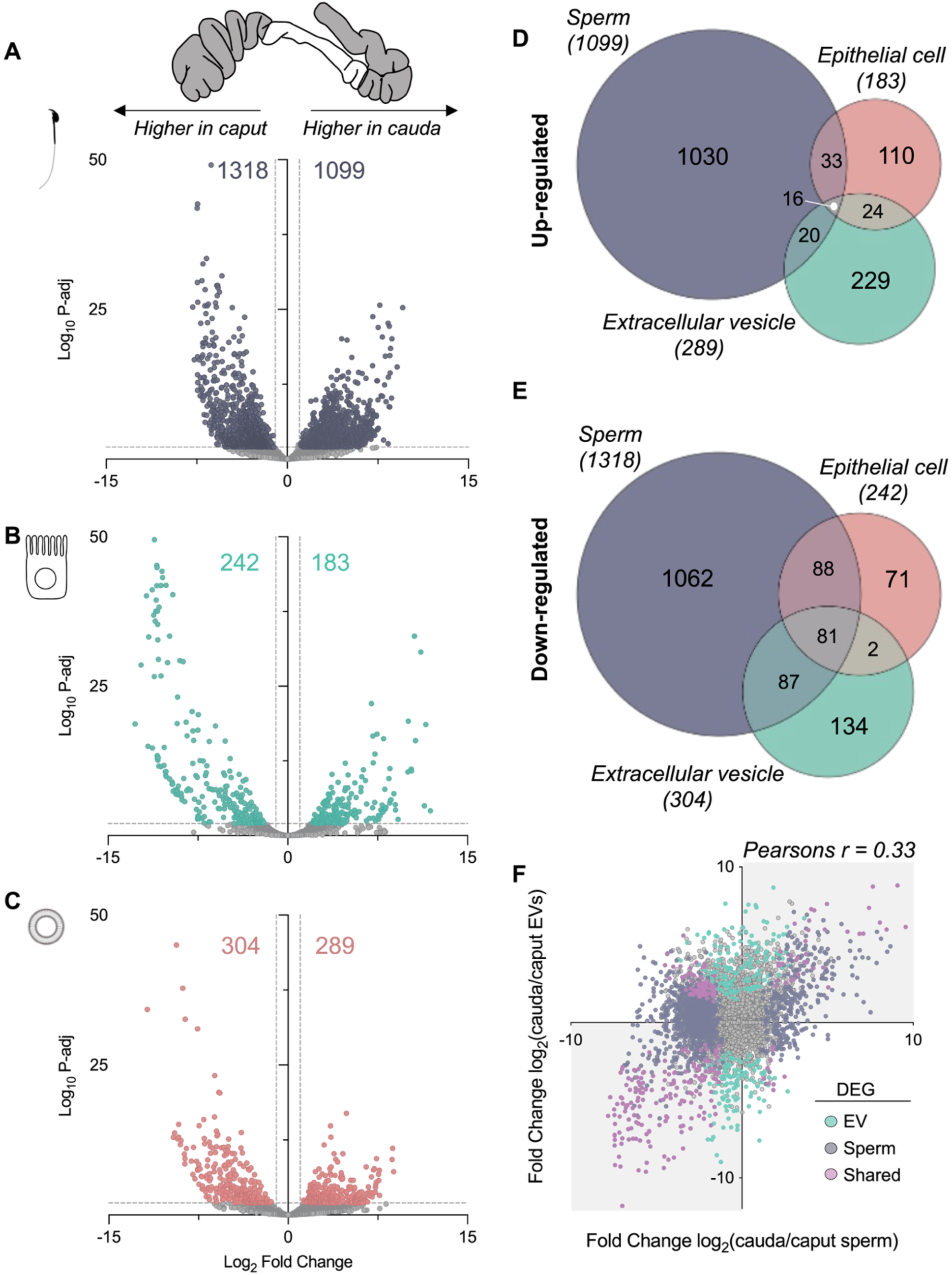
The mRNA profile of sperm, epithelial cells and epididymosomes is modulated along the epididymis. A-C) Volcano plots depicting log_2_ fold change and log_10_ adjusted *P*-value of identified genes in populations of A) sperm, B) epithelial cells and C) extracellular vesicles (EVs) isolated from the caput and cauda epididymis. Colored dots indicate differentially abundant genes as determined using DESeq2 in R. Significance threshold: Fold change ± > 2 and adjusted *P*-value of ≤ 0.01. D-E) Venn diagrams illustrating the overlap of D) up-regulated and E) down-regulated genes (DEGs) in sperm, epithelial cells and extracellular vesicles when cauda samples were compared to caput samples. F) Correlation plot of cauda/caput log_2_ fold change of mRNA expression in sperm (x-axis) and EVs (y-axis). Colored dots highlight DEGs in sperm (purple), EVs (green) and those genes altered in both samples (pink) when cauda samples are compared to caput.

### Extracellular vesicles convey mRNAs to sperm following in vitro co-incubation

Based on our findings that mRNA accumulation in EVs and sperm exhibit strikingly similar dynamics during epididymal transit and, importantly, the previous demonstration of small RNA and protein transfer between sperm and EVs, we performed *in vitro* co-incubation experiments of caput sperm and cauda EVs to determine if EVs transfer mRNAs to epididymal sperm (Sharma et al., 2018, Reilly et al., 2016, Nixon et al., 2019). Following co-incubation and thorough washing, RNA was extracted from populations of sperm co-incubated with EVs and naïve sperm with control co-incubation (Fig. S2 A-B). This analysis revealed the increased abundance of mRNAs from 319 genes in sperm co-incubated with EVs and 217 genes with decreased abundance (Fig. 3A). Intriguingly, comparing the 319 up-regulated genes to the list of cauda sperm specific transcripts, mRNAs expressed at ≥ 5 TPM in cauda but not caput sperm, we identified 77 genes shared between these groups (Fig. 3B). Owing to the technical challenges in mimicking epididymal sperm exposure to EVs (2 h in vitro co-incubation compared to 7-10 days of exposure *in vivo*), to quantitate more subtle alterations in sperm mRNAs we sorted genes increased in abundance in cauda sperm compared to caput sperm into bins grouped by fold enrichment in cauda sperm compared to caput sperm and examined the fold change of these groups between control caput sperm and caput sperm incubated with EVs (Fig. 3C). Plotting cumulative distribution illustrates the genes with greatest increase from caput to cauda sperm naturally (*in vivo*) demonstrated increased abundance in populations of caput sperm incubated with EVs compared to caput sperm mock incubated controls. In particular, the transcripts of 32 genes demonstrate an increased abundance when caput sperm are exposed to EVs (analogous to cauda sperm) compared to control incubated caput sperm, including *Sf1*, *Ubtd1* and *Poc1b*, which are also abundant in cauda epididymosomes (Fig. 3D-E).

**Figure 3:**
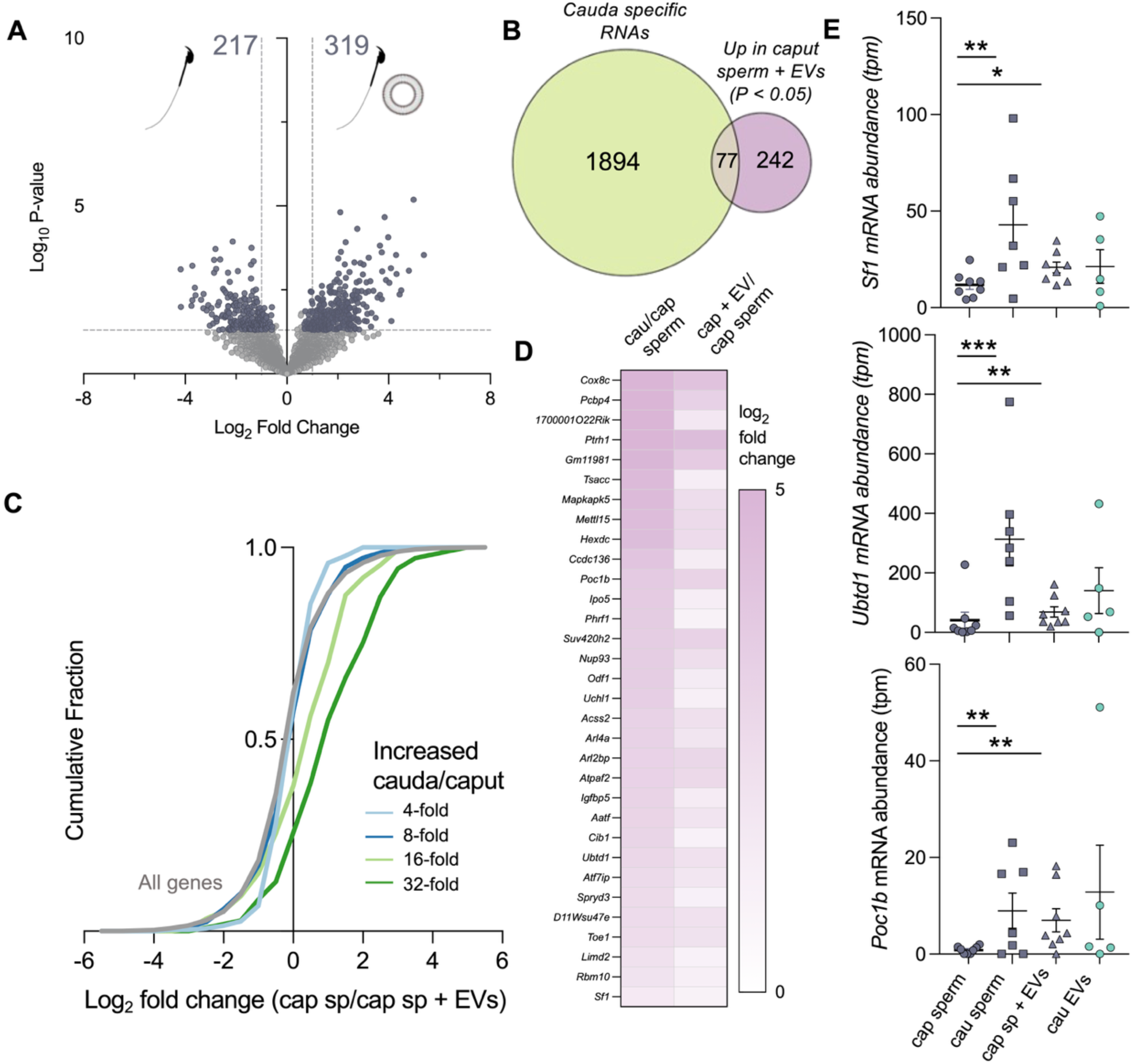
Epididymal EVs convey mRNAs to sperm following in vitro co-incubation. A) Volcano plot depicting the RNA changes between caput sperm (mock incubation) and caput sperm co-incubated with cauda extracellular vesicles (EVs) in vitro. Colored dots indicate differentially expressed genes. Threshold for determining differentially expressed genes was fold-change of ± 1.5 and *P*-value σ; 0.05. B) Venn diagram depicting cauda specific genes (not detected in caput sperm and >10 average transcript per million (TPM) in cauda sperm) and up-regulated RNAs in caput sperm co-incubated with EVs compared to caput sperm alone. C) Cumulative distribution plot (CDF) of log2 fold change between caput sperm only and caput sperm incubated with EVs for all genes (gray line) and genes that are significantly increased in cauda sperm compared to caput sperm (green and blue lines). Increased genes are separated into four bins based on cauda sperm/caput sperm fold change data from Fig. 2A. D) Heatmap illustrating the log_2_ fold change of the 32 genes significantly increased, both in cauda sperm and caput sperm incubated with EVs, compared to caput sperm alone. E) mRNA abundance (TPM) of *Sf1*, *Ubtd1* and *Poc1b* in caput and cauda sperm, caput sperm post-co-incubation experiment and cauda EVs. Each dot indicates an individual biological replicate, and statistical analysis was performed to determine differences between caput and cauda sperm and caput sperm with and without co-incubation of EVs. Asterisks indicate level of significance, where * = *P*-value < 0.05, ** = *P*-value < 0.01 and *** = *P*-value < 0.001.

### mRNAs are protected in extracellular vesicles

The enrichment of EVs using the protocol in this study does not discount the possibility of mRNA captured arising from unprotected extracellular mRNA that co-elutes with EVs. Hence, to discriminate EV encapsulated RNA from extracellular RNA not associated with EVs we treated EVs with RNase A, to remove any free RNA, using previously published methods (Fig. 4A)(Choy et al., 2022). Comparison of the detected RNAs in populations of EVs incubated in the presence of vehicle (mock) or RNases revealed the majority of mRNAs in both caput and cauda EVs were detected in both mock and RNase treated samples (Fig. 4B). Examining the average expression of RNAs within each of these samples showed an enrichment of mRNAs following treatment in populations of caput EVs, while cauda EV preparations treated with RNases demonstrated similar abundance to untreated (Fig. 4C). While not impacting vesicle integrity or morphology (Fig. S2B), RNase treatment did lead to the depletion of transcripts, suggesting their presence outside of the EVs. Ultimately however, this finding demonstrates that the mRNA sequenced in our EV preparations is derived from RNA encapsulated within EVs. Importantly, this includes many mRNAs we showed to be differentially regulated within EVs from the caput epididymis compared to the cauda epididymis (colored dots, Fig. 4C).

**Figure 4:**
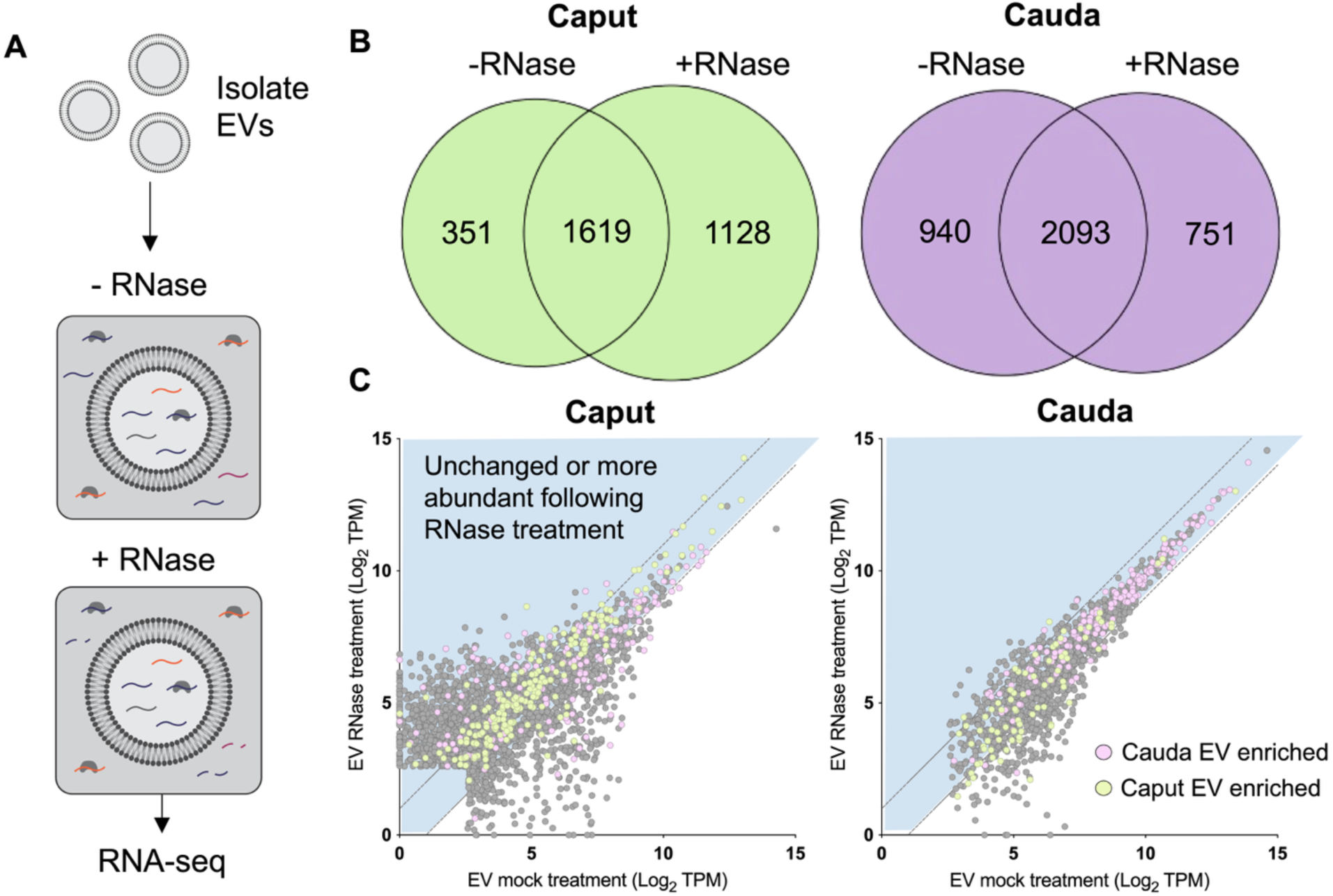
Epididymal mRNAs are protected in extracellular vesicles. A) Preparations of EVs isolated from the caput and cauda epididymis were split into equal aliquots and one sample was treated with RNase A and the other mock incubated without the addition of RNase. RNA was extracted following treatment and libraries generated for mRNA-seq. B) Venn diagrams illustrating the overlap of detected transcripts between mock and RNase treated preparations. C) Scatter plots of average log_2_ TPM for mock EV treatments (x-axis) versus RNase treated EVs (y-axis). Colored dots highlight mRNAs shown to be significantly altered in EVs between the caput and cauda epididymis. Purple indicates mRNAs enriched in cauda EVs and green indicates mRNAs enriched in caput EVs. The genes that lie in the blue shaded portion of the scatter plot highlight those that are either unchanged or more abundant in EV preparations following RNase treatment.

### Sperm transcripts not detected in the egg are found in the zygote

In alignment with published literature, we confirm that sperm carry mRNAs, however, whether these transcripts are delivered to the egg and function following fertilization remains unclear. To investigate this, we compared gene expression profiles of MII eggs and 1-cell embryos (zygotes 4.5 h post-fertilization; Fig. 5A, S3A). Using Venn diagrams to compare RNA profiles (detected genes are those with an average abundance of >5 TPM) from sperm, eggs and zygotes revealed most genes were shared among the three groups. Further, the egg and zygote shared the second highest number of expressed mRNAs from specific genes (Fig. 5B). Notwithstanding this, each cell type had a subset of unique mRNAs, with sperm displaying the greatest number of unique mRNAs. Of note, we identified 368 genes that were detected in both sperm and zygotes (>5 TPM average) but not detected in the egg. Among these were *Git1*, *Crcp, Cabyr* and *Ndufb7*. Further differential expression analysis revealed that 61.4% (226 genes) of these candidates were statistically more abundant in zygotes compared to eggs (Fig. 5B). Moreover, 42 of these genes were enriched in cauda sperm compared to caput sperm and 14 genes were also transferred to caput sperm by EVs in vitro (Fig. 3). Notably, 3 genes (*Toe1*, *D11Wsu47*e, and *Hexdc*) overlapped across the three of these categories (Fig. 5C), highlighting mRNAs transferred from the soma to sperm during epididymal transit and then to the zygote during fertilization (Fig. 5C).

**Figure 5:**
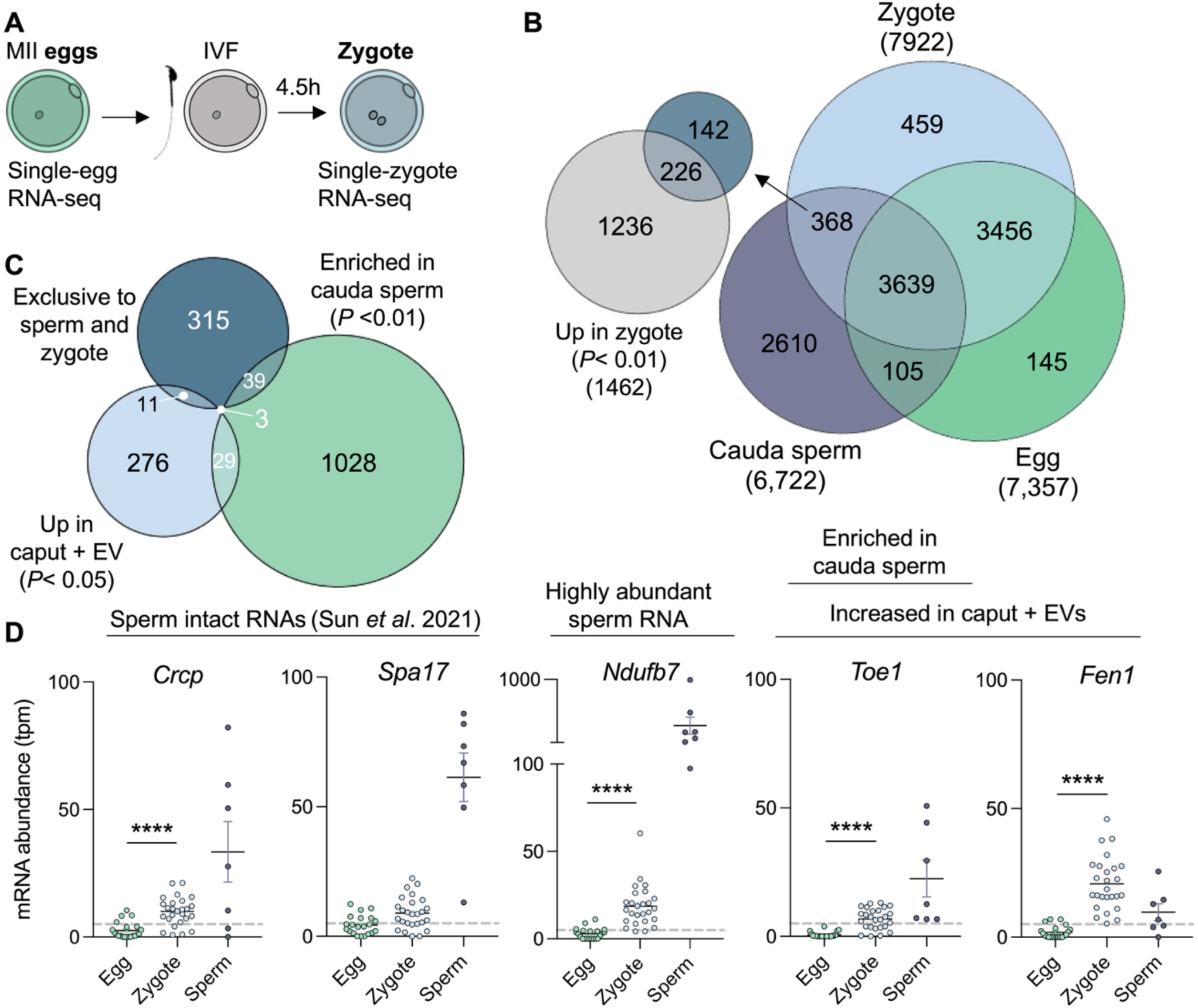
RNAs harbored by sperm are detected in fertilized eggs. A) Experimental schematic for collection of MII mouse eggs and fertilized eggs (zygotes). B) Venn diagram illustrating the overlap of detected genes in mature sperm, MII eggs and zygotes. Each gene list was filtered to generate a list of expressed genes (see Methods). Inset smaller Venn diagram illustrates the number of shared genes between those exclusively detected in sperm and zygotes and genes differentially expressed between egg and zygote (*P* < 0.01). C) Overlap of genes exclusively detected in sperm and zygotes, genes enriched in cauda sperm compared to caput sperm and those identified to be significantly increased in abundance in caput sperm following co-incubation with cauda EVs. D) Abundance of individual RNA transcripts in egg, zygote and sperm datasets. Individual genes were selected based on the presence in additional gene lists, including sperm intact RNA list (Sun et al., 2021), highly abundant cauda sperm mRNAs (from Fig. 2A) and increased abundance in caput sperm co-incubated with EVs (from Fig. 3A). Each dot represents a data point from a single egg, zygote or sperm replicate. Horizontal bar represents the mean and error bars depict standard error of the mean. Dotted line represents expression threshold of 5 TPM.

mRNAs in sperm have been thought to be remnant degradation products of spermatogenic and spermiogenic gene expression. While our mRNA-seq approach cannot rule out that the mRNAs we identify in sperm are mRNA-fragments, this is unlikely because our approach uses polyA-dependent cloning, thus only poly-adenylated RNAs will be sequenced. To further address this, we cross-referenced our results with a published dataset of ‘sperm intact RNAs’ (spiRNAs), confirming 803 spiRNAs in our cauda sperm dataset. Of these, 38 were uniquely present in sperm and zygotes but absent from eggs, including *Crcp* and *Spa17*, reinforcing that sperm can deliver intact mRNAs to the egg during fertilization (Fig. 5D).

### Mouse sperm mRNAs are conserved and expressed in human sperm

Following the identification of a population of mRNAs harbored in mouse sperm that are acquired by sperm as they transit the epididymis and delivered to the egg, we sought to examine the conservation of these sperm mRNAs in human sperm. To determine the human orthologs of mouse mRNAs we used Ensembl’s Bioconductor BiomaRt tool for R. This analysis identified human orthologs for 76% of the mouse transcriptome (Fig. 6A). Interestingly, the percentage of mouse mRNAs with human orthologs increased when this analysis focused on the subset of mRNAs transmitted to the zygote or epididymal acquired, with 96.6% and 95.0%, conserved, respectively (Fig. 6A). When comparing the abundance of all detected mRNAs in sperm across both species, we observed a significant correlation of expression (Pearson’s correlation coefficient = 0.132; Fig 6B), indicating that human and mouse sperm express overlapping mRNAs. Moreover, the sperm mRNAs acquired by EV co-incubation and those transmitted to the egg were detected and abundant in human sperm (Fig 6B). These findings suggest that sperm mRNA epididymal dynamics and the transmission of mRNAs to the zygote during fertilization may be conserved in mice and humans.

**Figure 6:**
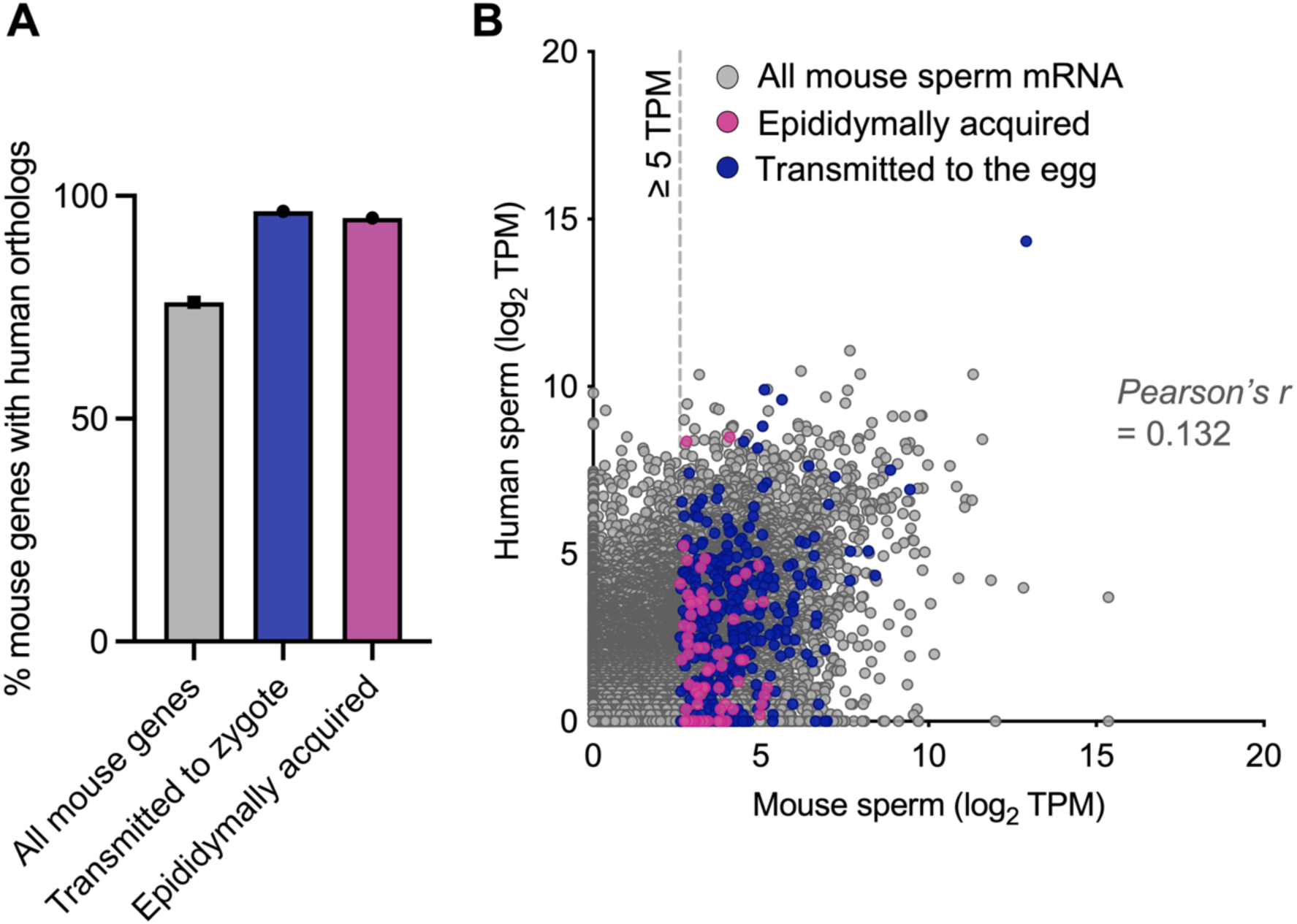
Conservation of mouse sperm mRNAs in human sperm. A) Bar plot illustrating the percentage of mouse sperm mRNAs with orthologs detected in human sperm. Columns depict all mouse sperm mRNAs (gray), mRNAs determined to be delivered to the egg (blue) and those acquired during epididymal transit (pink). B) Scatter plot illustrating the average mRNA abundance (log_2_ transcripts per million: TPM) of the 18,056 mouse sperm mRNAs that had human identified orthologs. Colored dots highlight mRNAs acquired by sperm during epididymal transit (pink) and transmitted to the egg at fertilization (blue).

### Sperm RNAs regulate postfertilization embryonic gene expression

Our data demonstrate the dynamic mRNA profile of sperm along the epididymis, the contribution of EVs to the acquisition of sperm mRNAs, and the delivery of mRNA to the egg. Additionally, total RNA extracted from mouse cauda sperm and introduced into parthenotes through microinjection has been shown to regulate ∼10% of genes in the early embryo (Conine et al., 2020). However, delineation of the impact of different sperm RNA fractions has not been explored. We next examined whether large RNAs present in sperm can modulate embryonic gene expression, similar to the established post-fertilization functions of sperm small RNAs (Trigg and Conine, 2024). To address this, we microinjected parthenotes with either control RNA (H3.3-GFP mRNA alone), total, or big RNA (>200 nucleotides) extracted from mature mouse cauda sperm. Parthenotes are chemically activated eggs that undergo development in the absence of a sperm cell, allowing for the unique detection of the function of sperm RNAs. Injected parthenotes were cultured and collected at the 4-cell and morula stage for single-embryo RNA-seq, along with biparental (sperm-fertilized) embryos generated via IVF with eggs from the same cohort of superovulated females used for parthenogenesis (Fig. 7A).

**Figure 7:**
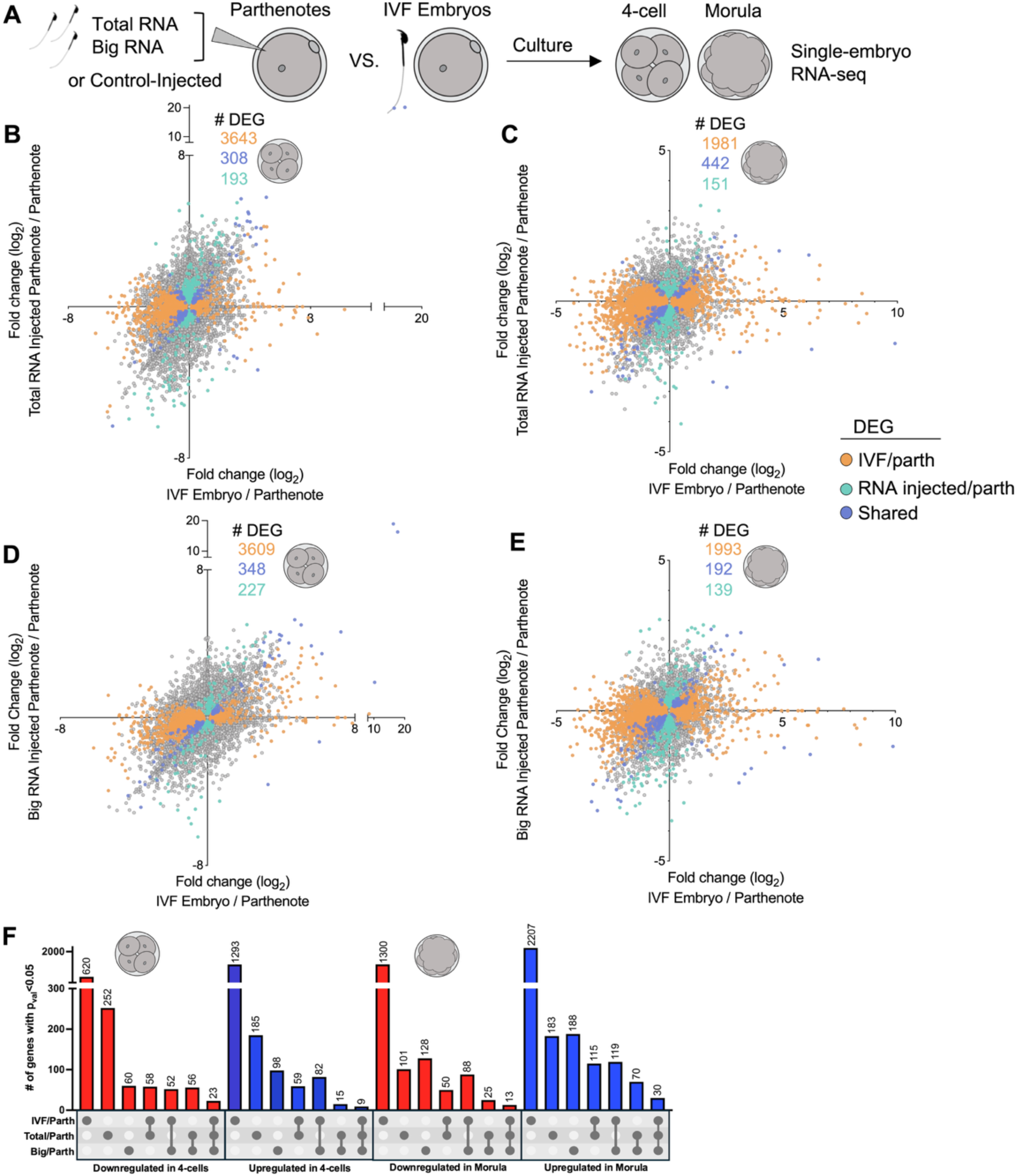
Sperm RNAs influence gene expression in parthenotes. A) Schematic of microinjection experiment involving the injection of sperm RNA into parthenotes and comparison to matched set of IVF generated embryos. B-E) Correlation plots comparing the log_2_ fold change of genes between IVF produced embryos and parthenotes (x-axis) to RNA injected parthenotes and control parthenotes. Data for (B) 4-cell and (C) morula for total RNA injections and comparisons for (D) 4-cell and (E) morula for big RNA (> 200 nucleotides) injections. Colored dots indicate differentially expressed genes (DEGs) for IVF embryos versus control parthenotes (orange), RNA injected parthenotes compared to control parthenotes (blue), and those that were similarly altered in both groups (purple). F) Upset plot depicting DEG sets in 4-cell and morula embryos for all comparisons. Colored bars indicate the fold change direction, where red indicates increased abundance compared to control parthenotes and blue indicates decreased abundance.

This analysis revealed large transcriptomic changes between control parthenotes, and IVF generated embryos, at both developmental stages, as expected. However, additionally, the introduction of either sperm total or big RNA led to the differential expression of genes in parthenotes at the 4-cell and morula stage, compared to control (Fig. 7B-E). The overlap of DEGs between total and big RNA injected parthenotes compared to controls was 72 and 95 for 4-cell and morula stage embryos, accordingly (Fig. S4A-B). Importantly, we identified 193 and 151 shared DEGs in parthenotes supplemented with total RNA and IVF embryos when compared to control parthenotes at the 4-cell and morula stage, respectively (Fig. 7B-C, blue dots), indicating that sperm RNA supplementation can program parthenote gene expression to be more embryo-like. Overall, the correlation of gene expression between sperm total RNA injections and embryos compared to control injected parthenotes was 0.16 in 4-cell and 0.08 in morula embryos, indicating wide-ranging alterations in gene expression modulated by sperm RNAs that mimic fertilization. Further, when sperm big RNA was injected, the correlation increased to 0.26 and 0.09, for 4-cell and morula, respectively. Indeed, 227 genes were equally altered in big RNA injected parthenotes and IVF embryos compared to control parthenotes at the 4-cell stage, while at the morula stage we identified 139 shared DEGs, demonstrating that for many genes the introduction of sperm RNA >200 nucleotides resulted in a parthenote’s transcriptional profile more closely resembling that of IVF derived embryos.

Comparison of DEGs across all datasets revealed 32 genes shared between IVF, total and big RNA injected parthenotes compared to control parthenotes at the 4-cell stage, while 43 genes were shared at the morula stage. Consistent with delivery of sperm small RNA (included in total RNA), total RNA-injected parthenotes displayed a reduced expression of a subset of genes previously identified to be regulated by sperm small RNAs during preimplantation embryonic development, which were not regulated when sperm big RNA was supplemented (Fig. S4C-D). However, notably, parthenotes injected with big RNA shared more DEGs with IVF embryos than total RNA injected parthenotes (Fig. 7F), indicating these RNAs can indeed modulate postfertilization embryonic gene expression.

## DISCUSSION

Sperm harbor a diverse profile of RNA, including coding and non-coding RNAs. The latter of these, specifically small non-coding RNAs, have been demonstrated to causally transmit environmentally modulated non-genetically inherited phenotypes to offspring (Sharma, 2019, Lee and Conine, 2022). Conversely, while mRNAs were among the first RNA reported in sperm, they have received comparatively little investigation. This is likely due, in part, to the incongruity between the relatively low yield of sperm RNA and the large amount of RNA stored in the egg, as well as the assumption that mRNAs detected in sperm are merely remnants of late-stage spermatogenesis. However, emerging evidence has begun to identify postfertilization roles for specific transcripts (Cannarella et al., 2024, Yao et al., 2010). Here, we sought to determine whether, like sperm proteins and small RNAs, mRNAs are dynamically modulated throughout epididymal transit and the role of extracellular vesicles (EVs) in facilitating this modulation. Our analysis revealed the dynamic nature of mRNAs isolated from sperm, epithelial cells and EVs along the epididymis, with sperm displaying the most significant change to their mRNA profile from the caput to cauda epididymis. Moreover, we confirm the transfer of mRNA via EVs *in vitro* and importantly demonstrate that >200 nucleotide RNAs harbored in sperm function post-fertilization to regulate embryonic gene expression.

### Epididymal contribution to the sperm mRNA profile

Our data reveal that sperm epididymal maturation is accompanied by modulation of the sperm mRNA profile of sperm. Interestingly, this modulation sees the loss of greater than 1,300 mRNAs in sperm from the caput epididymis to the cauda (Fig. 2A). While not yet experimental validated, the loss of macromolecules from sperm has been attributed to the shedding of the residual cytoplasm (in the form of the cytoplasmic droplet) that occurs early in the epididymis (Cooper, 2011). Conversely, approximately 1,000 mRNAs increase in abundance in cauda sperm compared to caput sperm. The predominance of mRNA loss over gain in sperm during epididymal transit mirrors observations from their proteomic cargo (Skerrett-Byrne et al., 2022). The main form of intercellular communication in the epididymis, responsible for delivering proteins and small RNAs to epididymal sperm, are EVs. Resulting from the transcriptionally and translationally silent state of epididymal sperm, this form of intercellular communication represents a mechanism that shapes the sperm macromolecular cargo facilitating sperm maturation. Indeed, we demonstrate here, that epididymal EVs also transfer mRNAs to sperm (Fig. 3A). Co-incubation of caput sperm with cauda EVs revealed the increased abundance of 319 mRNAs, 77 of which are exclusively expressed in cauda sperm. Moreover, interestingly 217 mRNAs were reduced in caput sperm co-incubated with EVs compared to control caput sperm. This observation suggests a loss of mRNA during in vitro co-incubation, potentially reflecting active molecular exchange. Although yet to be experimentally validated, EVs have been proposed to facilitate the bidirectional transfer of molecules to refine the sperm macromolecular cargo (Zhou et al., 2019). While these changes do not encompass the breadth of alterations that occur from caput to cauda in vivo, they demonstrate the ability of EVs to transfer big RNA species to sperm. The additional time spent in contact with EVs in vivo (7-10 days) as well as other modes of intracellular communication must cooperatively shape the sperm mRNA profile. Indeed, nanotubules, the delivery of RNA via RNA-binding proteins, and the cytoplasmic droplet are all mechanisms of refining the sperm macromolecular cargo that could contribute to the sperm mRNA profile modulation (Battistone et al., 2019, Arroyo et al., 2011, Wang et al., 2023).

Several highly abundant mRNAs in cauda sperm are well-known spermatogenic transcripts that encode proteins required for sperm to function and thus important for male fertility. For example, *Spa17* encodes a protein (SP17) involved in sperm-zona pellucida interaction, while *Smcp* encodes a mitochondria associated cysteine rich protein that has been shown to enhance sperm motility (Nayernia et al., 2002, Brewis and Wong, 1999). The presence of such transcripts at high abundance in mature sperm suggests that many transcripts are indeed remnants from spermatogenic gene expression during testicular development. However, we found that *Smcp* is significantly increased in abundance in cauda sperm compared to caput sperm, an expression pattern also seen in epididymal tissue (Robertson et al., 2020). Thus, genes typically expressed in the testis also demonstrate abundance levels in epididymal sperm that suggest an epididymal contribution. Interestingly, while abundant in cauda EVs, *Smcp* is not detected in either caput or cauda epididymal epithelial cell preparations. Owing to prolonged incubation of single cell suspensions during epididymal epithelial cell preparations it is unlikely these cell populations would contain clear cells, a population of cells that express *Smcp* and that produce their own population of epididymal EVs (Battistone et al., 2019).

Our analysis identified many mRNAs that displayed concomitant increase in sperm and EVs from the caput to cauda epididymis (Fig. 2F), suggesting the involvement of EVs in delivering these mRNAs to sperm. Notable examples of such mRNAs include, *Akr1b7*, *Azin2*, and *Gpx3*, which encode proteins that are broadly involved in redox regulation and metabolic support for sperm maturation (Lambertos et al., 2018, Schwaab et al., 1998). Moreover, one of the mRNAs most significantly increased in cauda sperm compared to caput sperm was *Crisp1*. This mRNA is not expressed in the testis, and its protein product CRISP1 is well-known to be secreted from the epididymis and involved in several key sperm functional maturation events (Trigg et al., 2025, Sulzyk et al., 2025, Hu et al., 2018). Here, we show abundant *Crisp1* mRNA levels in sperm, EVs and epithelial cells from the cauda epididymis, but low expression in the caput epididymis. CRISP1 protein is also increased in abundance in cauda sperm, compared to caput sperm, with its acquisition important for sperm functionality (Skerrett-Byrne et al., 2022, Hu et al., 2018). This increase in CRISP1 protein levels alongside elevated *Crisp1* mRNA in the sperm cell is surprising given the translationally silent state of spermatozoa. Moreover, given the inability of sperm to synthesize new proteins, a suggested role for an epididymal contribution to the sperm mRNA profile could be for delivery to the egg at fertilization for translation in the zygote (Fang et al., 2014). However, for the mRNAs mentioned thus far, this contrasts with the current known functions of their protein products in events surrounding sperm functional maturation, such as motility and sperm-egg interaction. Therefore, further research to understand the functional significance of this subset of sperm related mRNAs accumulated during epididymal transit should be explored.

### Sperm deliver mRNAs to the zygote

Owing to the lack of translational machinery within epididymal sperm, the fate of these mRNAs carried by sperm could be post-fertilization functions upon delivery to the egg cytoplasm, through either translation of the mRNAs or non-coding mechanisms. Indeed, we identified 368 mRNAs in the zygote that were also abundant in sperm samples but not detected in the egg. Importantly, a portion of these mRNAs were also enriched in cauda sperm and delivered to caput sperm through EV co-incubation (Fig. 5C). Thus, demonstrating the modulation of these mRNAs in the epididymis prior to delivery to the zygote. Of the mRNAs exclusive to sperm and zygotes was *Cabyr,* the calcium-binding tyrosine phosphorylation-regulated protein encoded transcript which has been previously shown to be transferred by sperm to the egg in humans and bulls (Johnson et al., 2015, Gross et al., 2019). The established role of *Cabyr* protein product is in modulating intercellular calcium levels during capacitation (Naaby-Hansen et al., 2002). Thus, its presence at the mRNA level in sperm and delivery to the zygote remain enigmatic. Although the post-fertilization roles of sperm mRNAs have had little investigation, our findings reveal an epididymal contribution to the sperm mRNA profile and subsequent delivery to the zygote signifying a functional role for sperm-borne mRNAs in the zygote.

While our RNA-seq approach selects poly(A) RNAs prior to fragmentation, therefore reducing the likelihood of sequencing mRNA fragments, the possibility of detecting mRNA fragments remains a limitation of the current study. To overcome this, we compared our sperm mRNAs to a publicly available list of sperm intact RNAs. Facilitated by long-read sequencing technologies a list of sperm intact RNAs (coined, spiRNAs) which represent full-length mRNA transcripts within both mouse and human sperm has been curated (Sun et al., 2021). Indeed, 3,440 spiRNAs were detected in mouse sperm, including 2,343 mRNAs and the remaining lncRNAs. Of the mRNAs identified in mature cauda sperm in our data, 803 were also identified spiRNAs, including *Cabyr*, *Smcp, Crisp1,* and *Spa17*. Importantly, for the clinical relevance of this work, we demonstrate the conservation of mouse sperm mRNAs in human sperm, suggesting potential conserved functions postfertilization or during sperm maturation (Fig. 6). Notably, a higher proportion of the mRNAs identified as being transmitted to the zygote had human orthologs compared to the overall mouse transcriptome.

### The functional role of sperm big RNAs

Microinjection of environmentally altered sperm RNAs, either isolated total, or fractionated small RNAs from sperm populations, or synthetic microRNAs (miRNAs), have been demonstrated to facilitate the inheritance of offspring phenotypes (Rodgers et al., 2015, Trigg et al., 2024, Sharma et al., 2016, Gapp et al., 2014, Tomar et al., 2024). Moreover, in line with previous reports, we demonstrate here that parthenogenetically activated eggs are a tractable model for isolating the functions of sperm RNAs in the absence of other sperm-derived factors (Conine et al., 2020). Indeed, we determined the ability of sperm RNA to regulate gene expression in the early embryo by microinjecting sperm purified total and big RNA (>200 nucleotides). Importantly, we confirm the downregulation of a subset of genes known to be regulated by sperm small RNAs, upon the introduction of sperm total RNA (Trigg and Conine, 2024). However, in line with the selection of RNAs greater than 200 nucleotides and therefore the removal of small RNAs, supplementation of sperm big RNA, did not lead to this downregulation (Fig. S4C-D). Further, we show for the first time, the rescue of 348 genes in 4-cell parthenotes mimics IVF embryos following the supplementation of sperm big RNAs (Fig. 7D). An interesting observation from our microinjection experiments is the higher correlation of gene expression in parthenotes injected with big RNA versus total RNA injected parthenotes compared to IVF at the 4-cell stage. Specifically, the loss of RNAs < 200 nucleotides led to greater correlation with IVF embryos than total sperm RNA injections which included 18-40 nucleotide small RNAs. This implies that big RNAs in sperm can influence post-fertilization embryonic gene expression at a level equal too, or greater than sperm small RNAs. The enrichment of RNAs > 200 nucleotides would include populations of mRNAs, circRNAs, and lncRNAs. Both circRNAs and lncRNAs have been shown to be altered in sperm in response to environmental cues. Moreover, the consequences of these altered sperm big RNAs has been linked to phenotypes in offspring, including metabolic and behavioral measures in paternal stress exposure models (Gapp et al., 2020, Hoffmann et al., 2024).

Collectively, our findings reveal a rich mRNA repertoire in mouse sperm influenced by the soma (epididymal epithelium), the transfer of these RNAs to the zygote at fertilization, and a previously underappreciated role of sperm RNAs, outside of small RNAs, in regulating embryonic gene expression. These findings reveal a wide array of new RNAs capable of functioning in RNA-mediated epigenetic inheritance. While we present evidence of post-fertilization functions for sperm RNAs >200 nucleotides in the regulation of gene expression in the early embryo, this does not preclude alternative functional roles in the male reproductive tract prior to ejaculation or in priming the female reproductive tract. However, our findings reveal a mechanism for the post-testicular modulation of sperm mRNAs by the soma (epididymis) which can be modulated by the environment or under stress conditions to transmit non-genetically inherited information to offspring. A comprehensive understanding of the role of sperm RNA role in inheritance requires extensive analysis of the diverse RNA classes present in sperm and the breadth of non-genetic information they encode.

## ACKNOWLEDMENTS

The authors gratefully acknowledge Rachel Bartlett, Liana Savarirayan, and Madeline Lamonica for mouse husbandry and talented rotation student Johnny Doherty for assistance with conducting experiments. We thank Teodora Orendovici and Stephen Mahoney of the Children’s Hospital of Philadelphia High Throughput Sequencing Core for technical assistance. We also thank Dr Biao Zuo from the TEM Core Facility at the University of Pennsylvania’s Electron Microscope Resources Lab. This work was funded by a Pew Biomedical Scholars Award to C.C.C., a Lalor Foundation Postdoctoral Fellowship (2022 and 2023) to N.A.T., a NICHD [F31 HD114433] to G.S.L, and a National Institutes of Health NIGMS T32 training grant (T32GM156697) to A.G.L.

## Supplementary Figures

**Figure S1:**
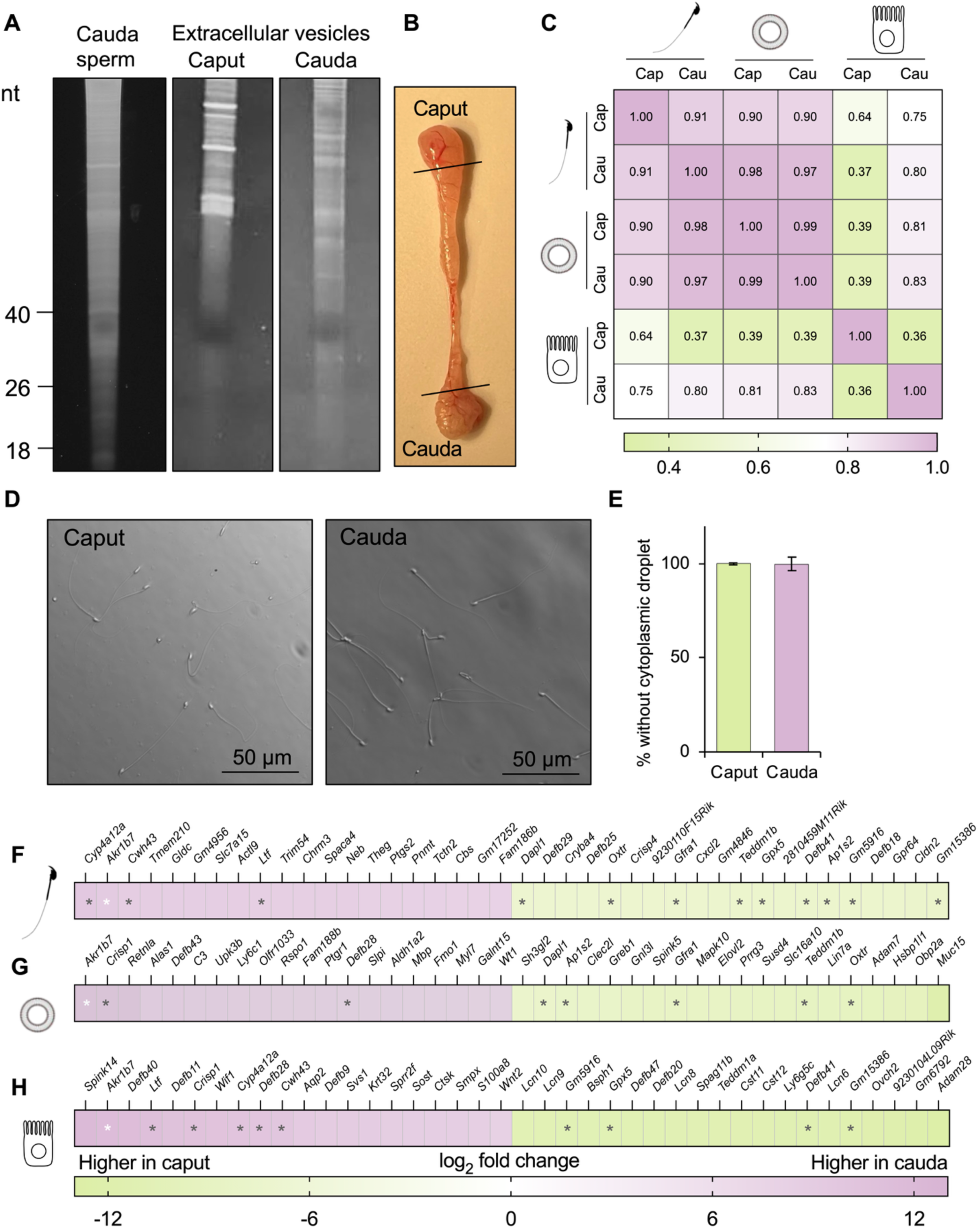
Epididymal sperm, extracellular vesicles and epithelial cell RNA-seq. A) Total RNA isolated from populations of cauda sperm and extracellular vesicles isolated from the caput, and cauda epididymis and separated using denaturing PAGE. Gel is stained with SYBR gold to stain nucleic acids. B) Image of a mouse epididymis highlighting the dissected epididymal segments representing the caput and cauda throughout this study. C) Pearsons correlation heatmap of sperm, epithelial cells, and extracellular vesicles from the caput and cauda epididymis. D) Phase microscopy images of purified caput and cauda sperm preparations following somatic cell lysis illustrating negligible somatic cell contamination. E) Graphical representation of the percentage of sperm devoid of cytoplasmic droplets in caput and cauda sperm preparations. F-H) Heat maps depicting the log_2_ fold change of the top 20 up-and 20 down-regulated mRNAs in cauda compared to caput in F) sperm, E) extracellular vesicles and H) epithelial cells. Asterisks indicate mRNAs that are also differentially expressed (DE) between caput and cauda in other sample types. Grey asterisks = DE in one other sample type, white asterisks = DE across all three sample types.

**Figure S2:**
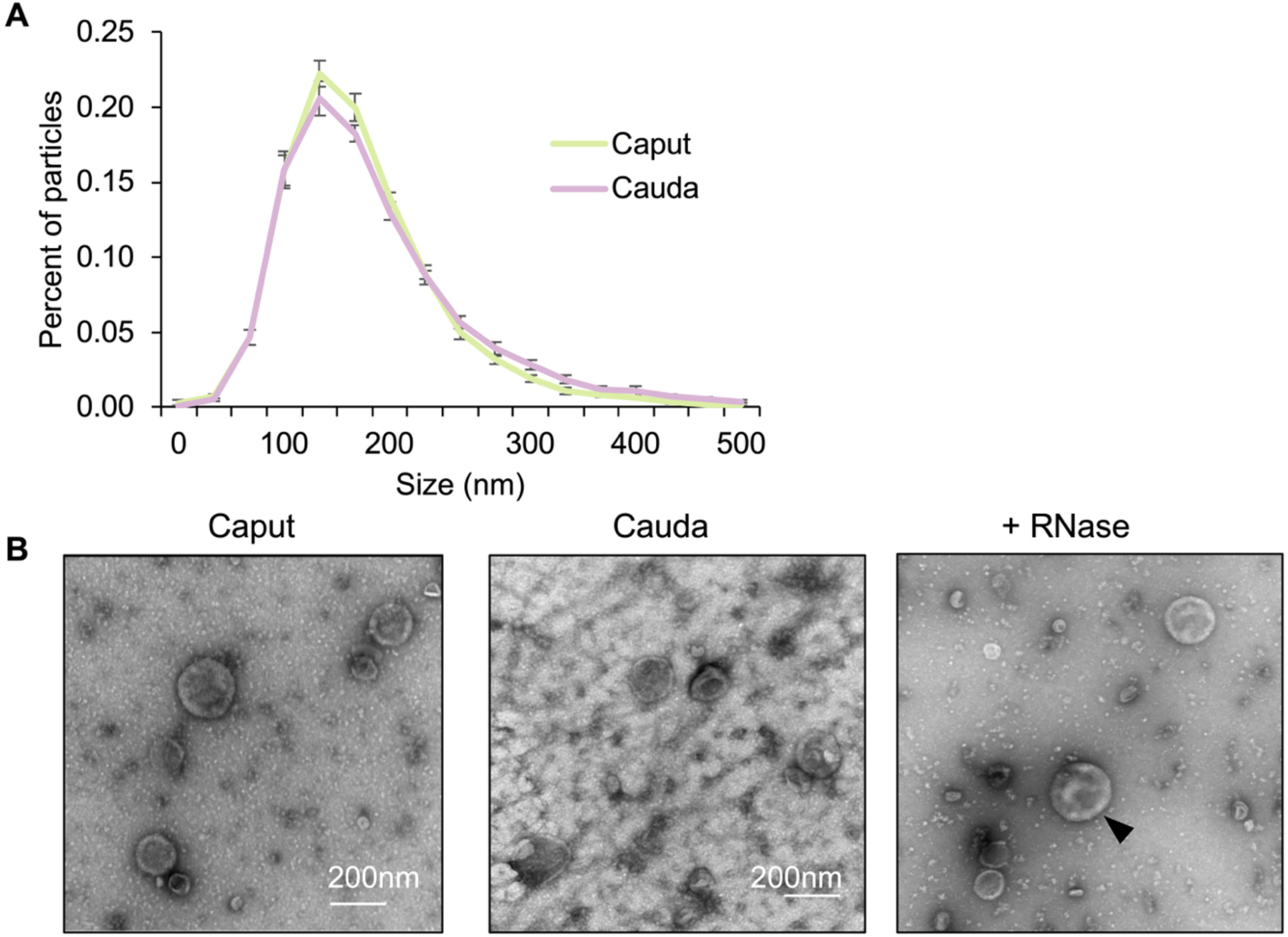
Characterization of epididymal extracellular vesicles. A) Vesicle size and distribution of caput and cauda extracellular vesicles (EVs). Data was obtained from ZetaView nanoparticle tracking software and is represented as mean with standard error of the mean. B) Negative stain electron microscopy images of EVs isolated from the caput and cauda epididymis and EVs post RNase treatment to visualize vesicle integrity following treatment. Scale bar = 200 nm.

**Figure S3:**
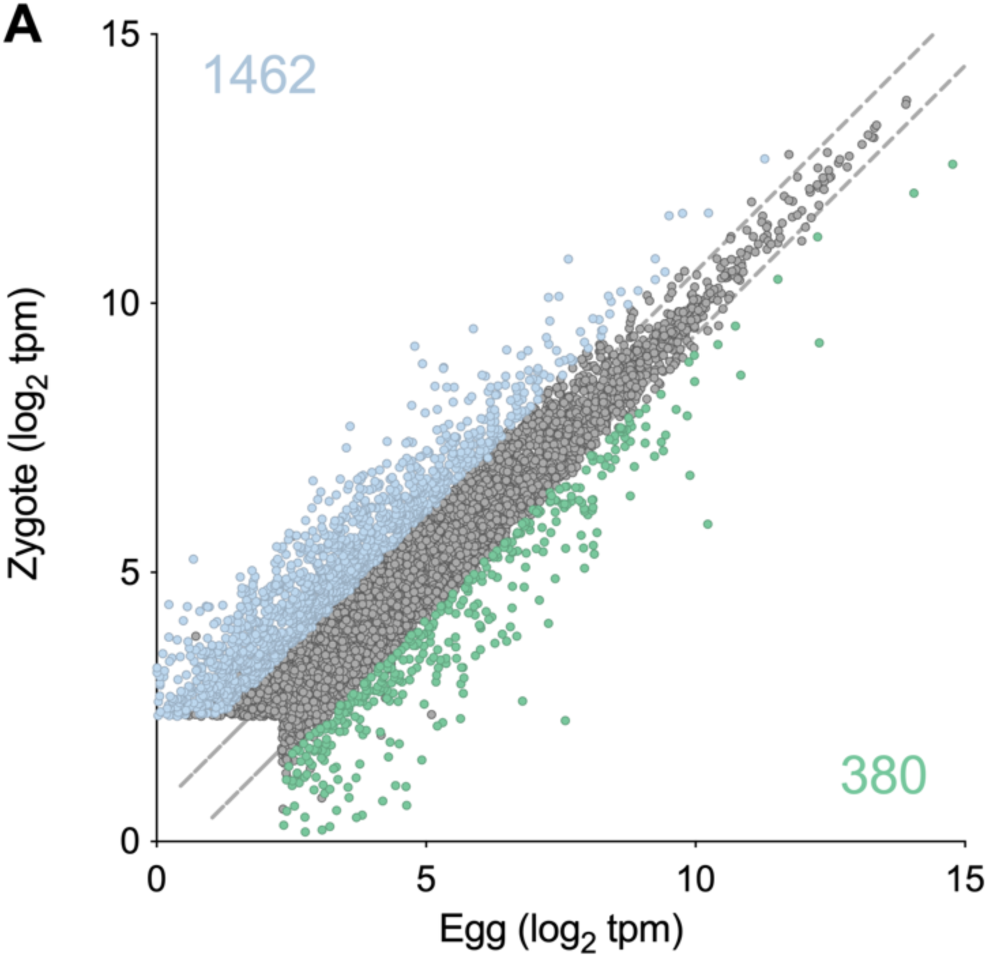
Transcriptomic alterations in mouse zygotes. A) Scatter plot illustrating average log_2_ gene expression (transcript per million; TPM) in mouse eggs (x-axis) and fertilized eggs (zygotes; y-axis). Colored dots indicate significantly altered genes; blue indicates genes higher in zygotes while green depicts higher expression in eggs. Threshold for differential expression was fold-change ± 2 and adjusted *P*-value of ≤ 0.01.

**Figure S4:**
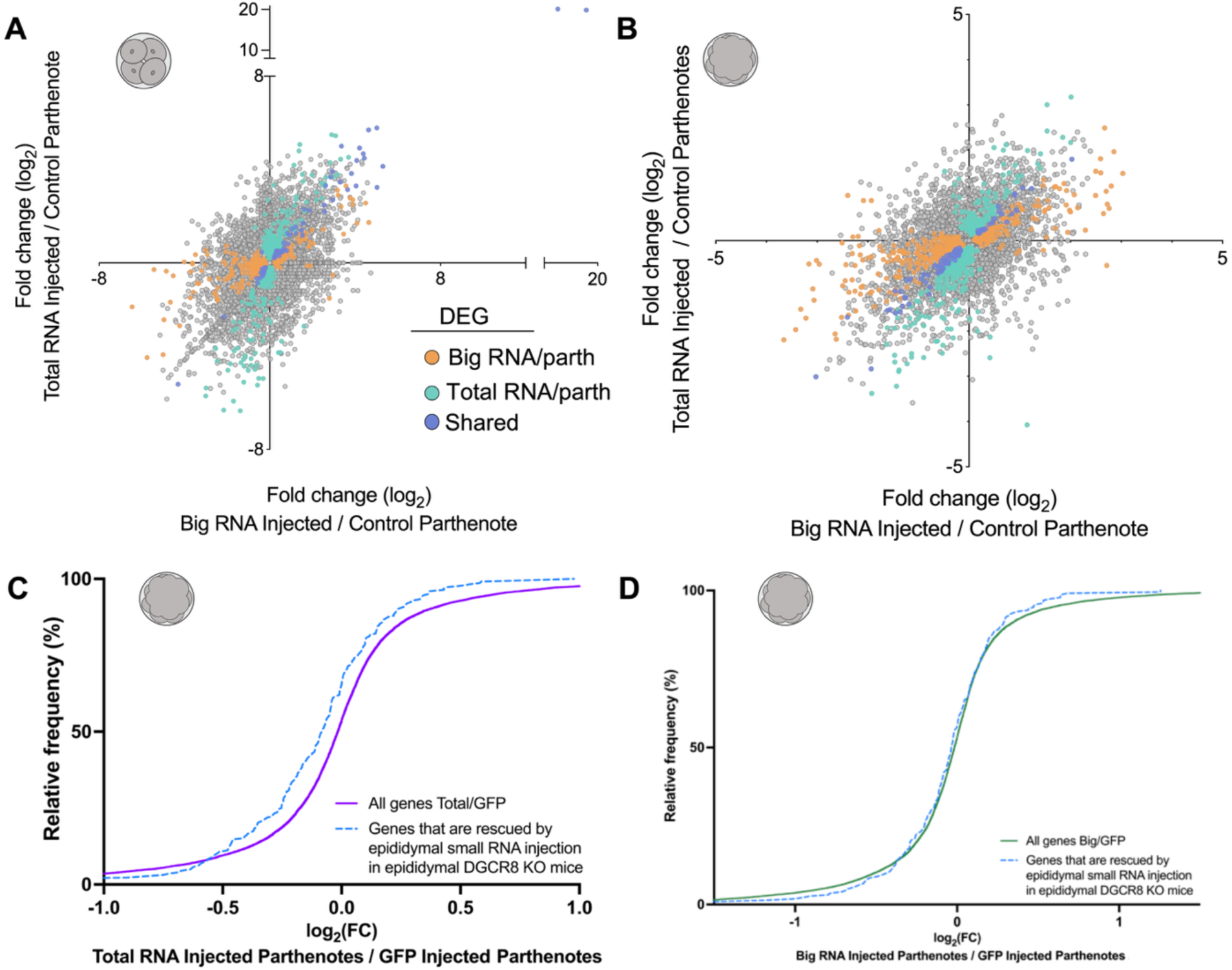
Sperm RNAs influence gene expression in parthenotes. A-B) Correlation plots comparing the log_2_ fold change of genes between big RNA and control injected parthenotes (x-axis) to total RNA and control injected parthenotes. Data for (A) 4-cell and (B) morula. Colored dots indicate differentially expressed genes (DEGs) for big RNA injected versus control parthenotes (orange), total RNA injected compared to control parthenotes (blue), and those that were similarly altered in both groups (purple). C-D) Cumulative distribution plot for log_2_ fold change in morula embryos for C) total and D) big RNA injected parthenotes compared to control parthenotes. Lines depicting all genes for total RNA injected versus control injected (purple) and big RNA injected versus control injected (green). Gene subsets are examined in all CDF plots with blue dotted lines demonstrating genes previously identified to be rescued in mutant embryos by injection of small RNA fraction, which would be included in the total RNA injection.

